## Supplemental information for "Gene-edited primary muscle stem cells rescue dysferlin-deficient muscular dystrophy"

### Supplementary figures and legends

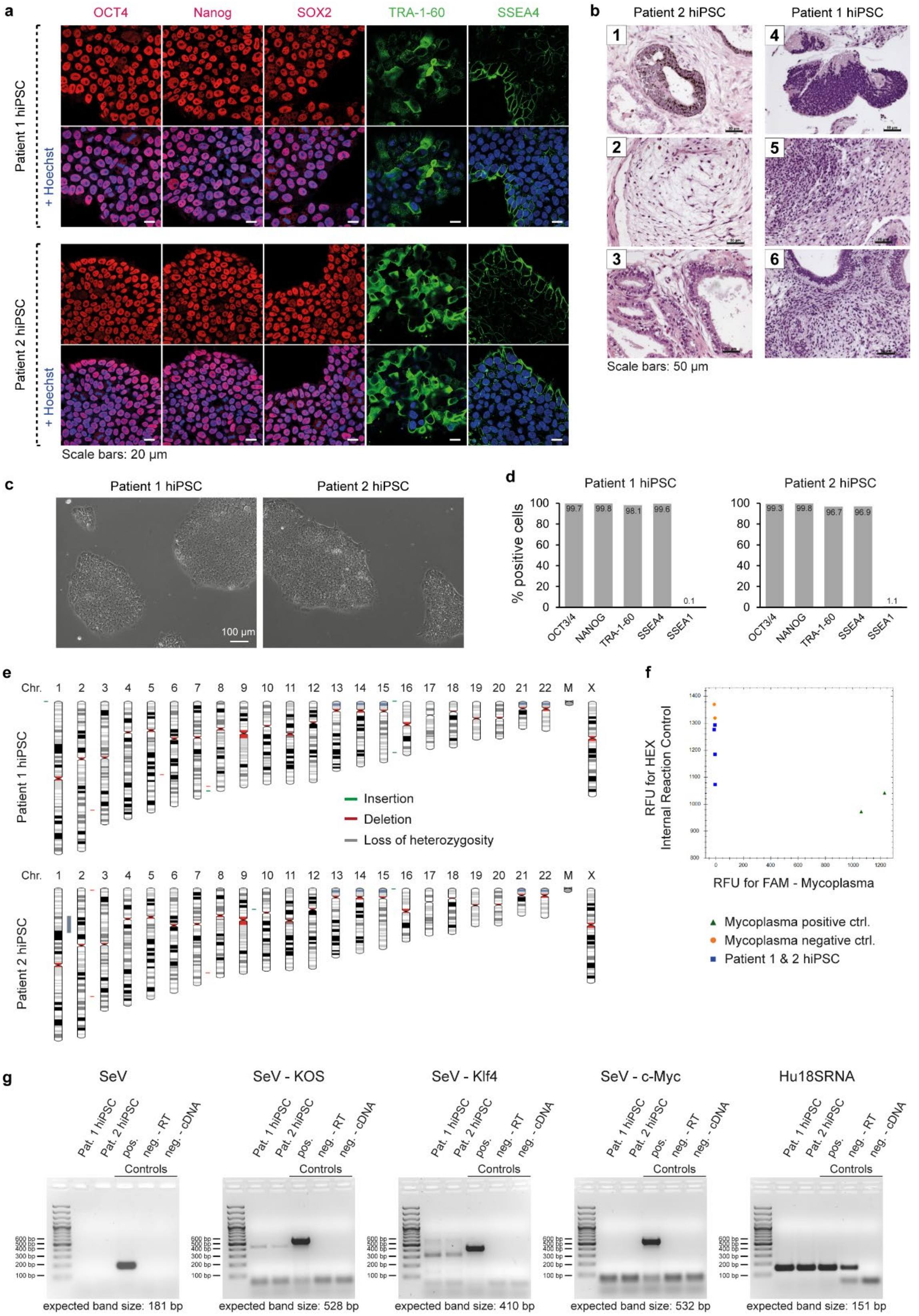


**Supplementary Fig. 1: Generation and characterization of patient hiPSC. a** hiPSC from two patients immunostained for pluripotency markers. **b** Histopathological analysis of patient hiPSC-derived teratomas with tissues from the three germ layers: (1) Primitive neurectoderm – beginning pseudorosette formation / Primitive ectoderm – pigment granula; (2) Primitive mesoderm – loose extracellular matrix;

(3) Endoderm – cuboidal epithelium and lining cyst-like structure; (4) Neuroectoderm – beginning rosette formation; (5) Mesoderm – immature mesenchyme and connective tissue; (6) Endoderm – isoprismatic to columnar epithelial cells. **c** Bright field microscopy images of hiPSC colonies cultivated in mTeSR1 medium and Matrigel coating. **d** Percentage of cells expressing pluripotency and differentiation markers analyzed by immunostaining and flow cytometry. **e** Virtual karyotype analysis. Regions of gain (duplications) are shown in green, regions of loss (deletions) are shown in red and regions of uniparental disomy (loss of heterozygosity) are shown in grey. Reportable are copy number changes greater than 0.4 Mb compared to the human reference genome and regions of loss of heterozygosity above 3 Mb. **f** Mycoplasma testing by RT-qPCR. X-axis shows relative fluorescence units (RFU) for FAM for Mycoplasma detection. Y-axis shows internal experimental control RFUs for HEX stain. **g** RT-PCR analysis of Sendai virus (SeV) genomes.

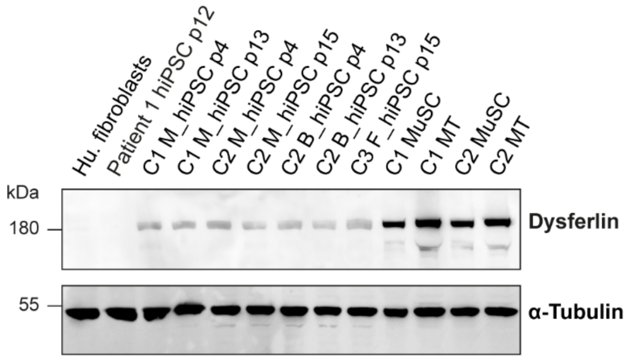


**Supplementary Fig. 2: Dysferlin is expressed in hiPSC.** Western blot analysis of dysferlin protein expression in hiPSC lines derived from MuSC (M_hiPSC), blood (B_hiPSC) or skin fibroblasts (F_hiPSC) from three controls (C1-C3) and from patient 1 at earlier and later passages after reprogramming (p4-p15). Human fibroblasts were used as negative control. Primary MuSC and myotubes (MT) from two controls were used as positive control. α-tubulin was used as loading control.

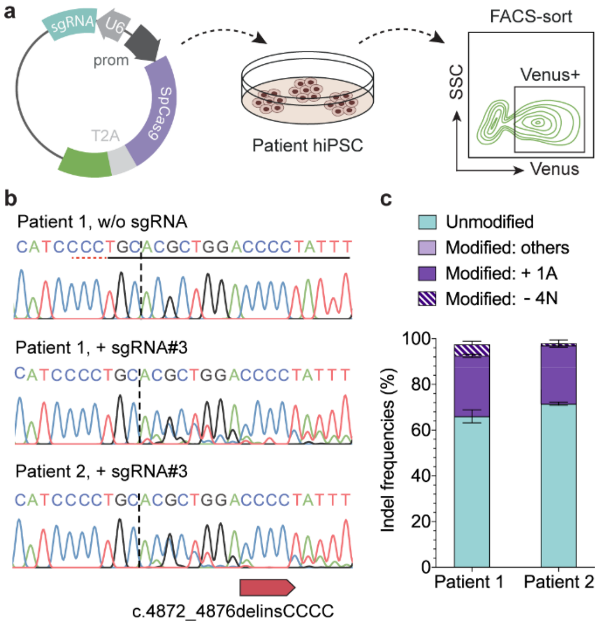


**Supplementary Fig. 3: Plasmid-based delivery of wild-type SpCas9 and sgRNA#3 efficiently reframes *DYSF* exon 44 in patient hiPSC. a** Schematic overview of experimental workflow. Patient hiPSC were transfected with a plasmid encoding for SpCas9 and a Venus fluorescence reporter, FACS-sorted, and processed for DNA analysis via Sanger sequencing. **b** Sanger sequencing chromatograms of edited (SpCas9, + sgRNA#3) hiPSC from both patients compared to unedited cells (SpCas9, w/o sgRNA). The protospacer and PAM sequences are underlined. The dotted vertical line indicates the expected DSB site. **c** Predicted indel frequencies based on chromatogram deconvolution analysis with ICE (Synthego) ($n$ = 2; mean ± SD).

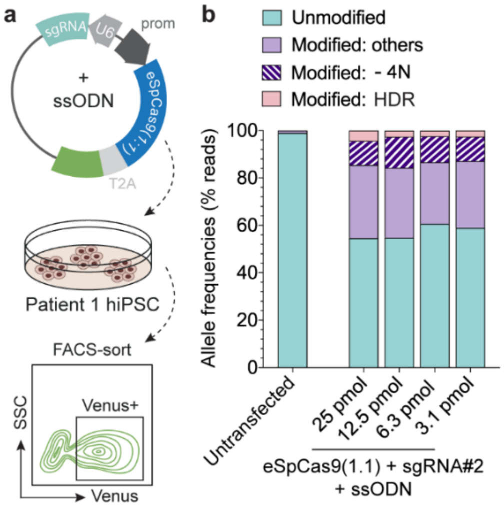


**Supplementary Fig. 4: CRISPR-Cas9 mediated HDR in patient hiPSC**. **a** Schematic overview of experimental workflow. Patient 1 hiPSC were transfected with a plasmid encoding for eSpCas9(1:1), sgRNA#2 and a Venus reporter, plus different concentrations of a donor template (in the form of a single-stranded oligodeoxynucleotide, ssODN) with the wild-type *DYSF* exon 44 sequence. Venus+ cells were selected by FACS-sorting and processed for DNA analysis via NGS. **b** Allele frequencies determined by NGS.

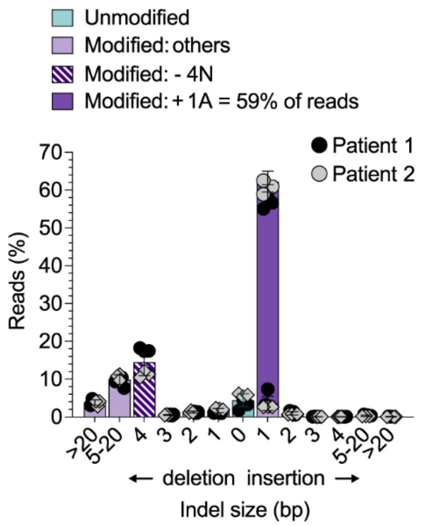


**Supplementary Fig. 5: Allele frequencies remain constant throughout cultivation time in edited patient MuSC. a** Frequency distribution of all indels in MuSC from the two patients at day 9 post-transfection with SpCas9 mRNA and sgRNA#3 (*n* = 3 replicates per patient, mean ± SD).

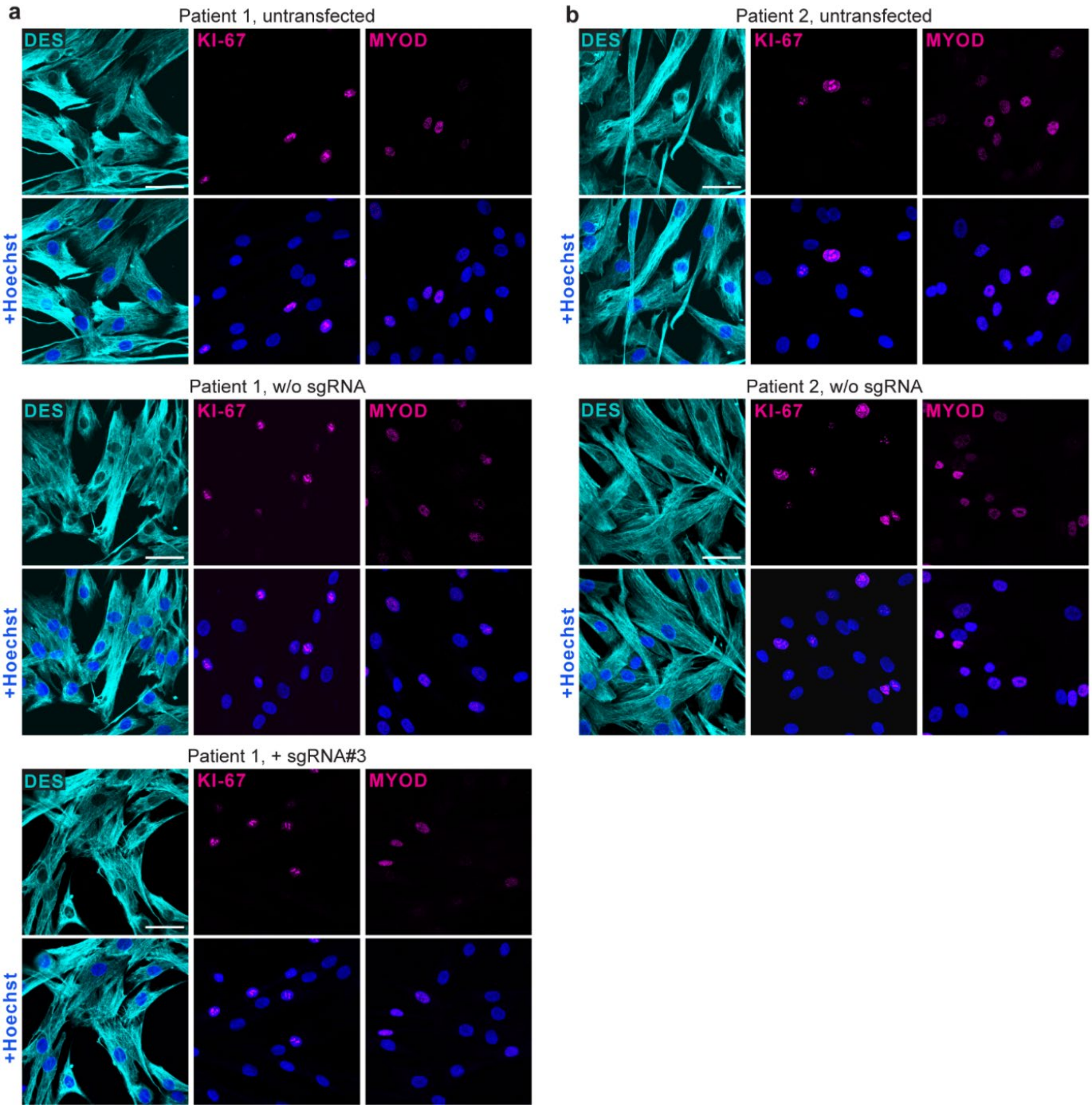


**Supplementary Fig. 6: Marker profiling of edited and unedited patient MuSC.** MuSC from patient 1 (**a**) and 2 (**b**) were stained for DES, PAX7 and MYOD 4 days after nucleofection with Cas9 mRNA, with or without sgRNA#3. The upper panels show untransfected cells from identical origin that were cultivated in parallel and processed for analysis at the same time. Blue: Hoechst. Scale bar: 50 µm.

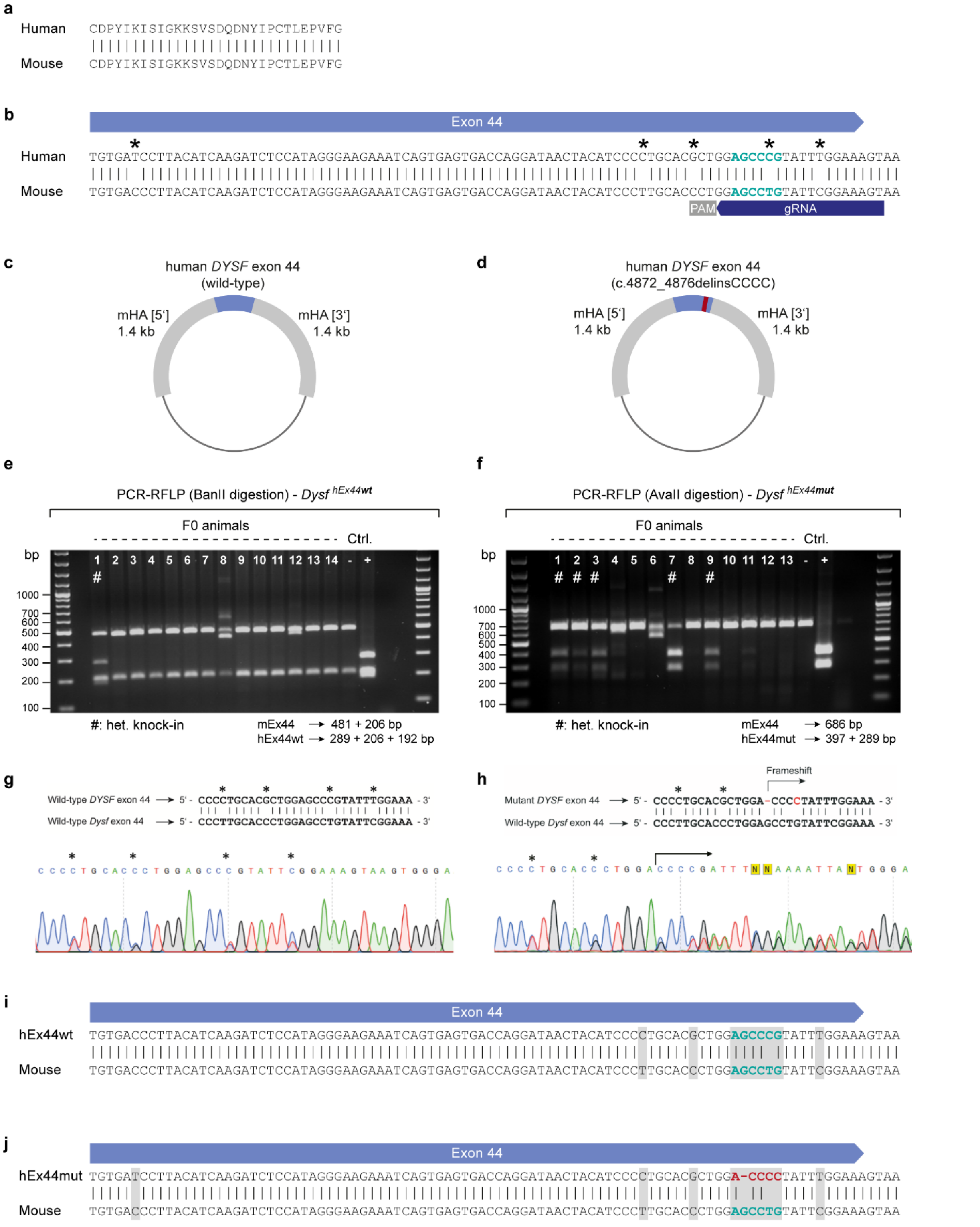


**Supplementary Fig. 7: Generation of a LGMD2B mouse model carrying the human *DYSF* exon 44 with the c.4872_4876delinsCCCC founder mutation and a corresponding control**. **a** Alignment of the amino acid sequence encoded by the human and murine wild-type *DYSF*/*Dysf* exon 44. **b** Nucleotide sequence alignment of the human and murine wild-type *DYSF*/*Dysf* exon 44. Mismatched positions are indicated with asterisks. The sequence corresponding to positions c.4872-c.4876 is highlighted in turquoise. The gRNA used to cut on the endogenous mouse exon 44 to generate the transgenic mouse lines is indicated. **c, d** Schematic overview of targeting vectors used to replace the murine exon 44 for the human wild-type (c) or mutant (d) *DYSF* exon 44. mHA: mouse homology arms. **e, f** genotyping strategy of hEx44wt (e) and hEx44mut (f) F0 animals by PCR plus restriction fragment length polymorphism (RFLP). Negative control: Genomic DNA from wild-type C57BL/6N mice. Positive control: Targeting vectors. **g, h** Chromatograms from heterozygous knock-in hEx44wt (g) and hEx44mut (h) F0 animals. **i, j** Nucleotide sequence alignment of the mouse wild-type exon 44 and the hEx44wt (i) and hEx44mut (j) newly generated alleles. The exchanged positions between the mouse genome and the targeting vectors are highlighted with grey squares.

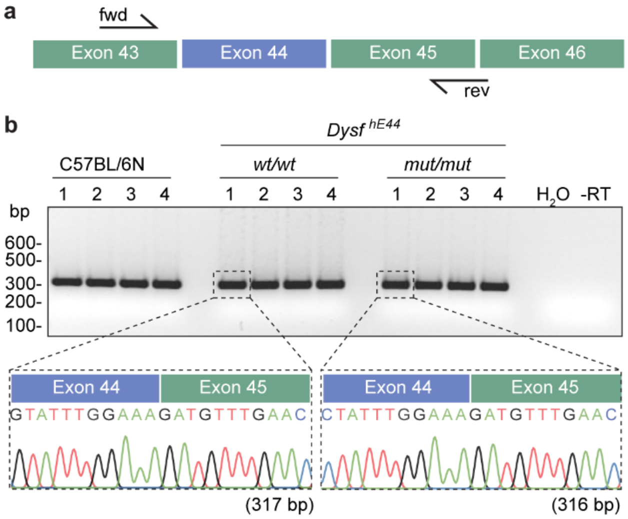


**Supplementary Fig. 8: mRNA analysis in homozygous *Dysf* exon 44 humanized mice. a** Primer design. **b** RT-PCR analysis of *Dysf* exon 43-46 in homozygous hEx44wt and hEx44mut mice. Wild-type C57BL/6N mice were used as control. The humanized *Dysf* exon 44 (wt or mut) is correctly spliced. The Sanger sequencing chromatograms below show the exon 44-45 junction in homozygous hEx44wt and hEx44mut mice.

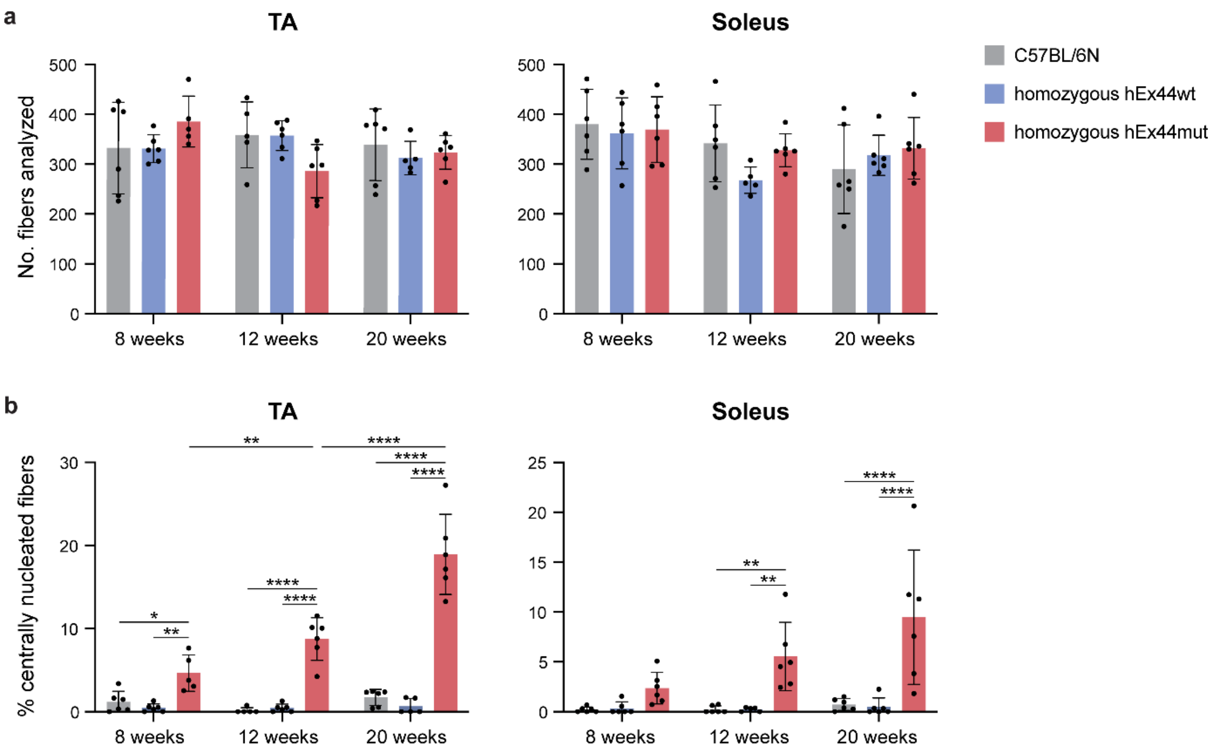


**Supplementary Fig. 9: Homozygous hEx44mut mice show a progressive dystrophic phenotype with onset at around 8 weeks of age.** Male homozygous hEx44wt and hEx44mut mice were phenotypically analyzed at age 8, 12 and 20 weeks using Gomori's trichrome histological stain and compared to male C57BL/6N wild-type mice. **a** Total number of fibers analyzed for TA (left) and Soleus (right) muscles for each mouse. **b** Percentage of centrally nucleated fibers in TA (left) and Soleus (right) muscles for each mouse. $n$ = 5-6 (mean ± SD); $p$ values were calculated using a 2-way ANOVA with Tukey's multiple comparisons test.

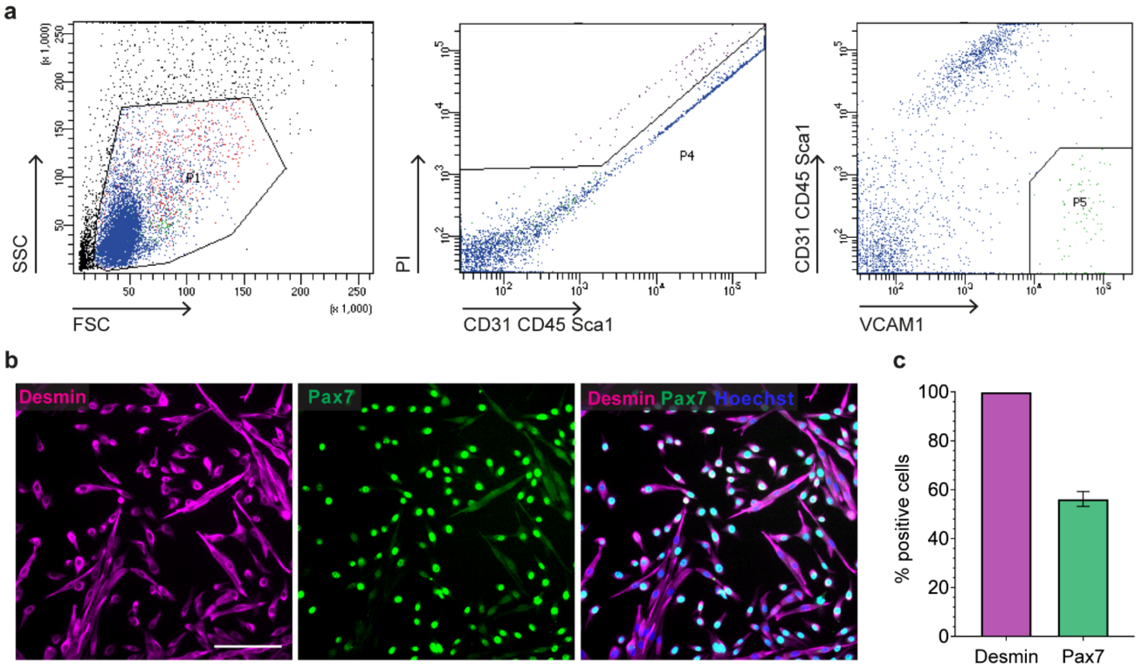


**Supplementary Fig. 10: MuSC isolation from *DYSF* exon 44 humanized mice. a** Gating strategy. Following tissue digestion and immunostaining, CD31-, CD45-, Sca1- (PE-conjugated primary antibodies) and VCAM1+ cells (Alexa Fluor 488-conjugated secondary antibody) were selected by FACS-sorting (P5 gate). Propidium iodide (PI) was used to exclude dead cells. **b** Immunostaining for myogenic markers Desmin and Pax7 one day after FACS-sorting. Scale bar: 200 µm. **c** Quantification of % cells positive for Desmin and Pax7 one day after FACS-sorting.

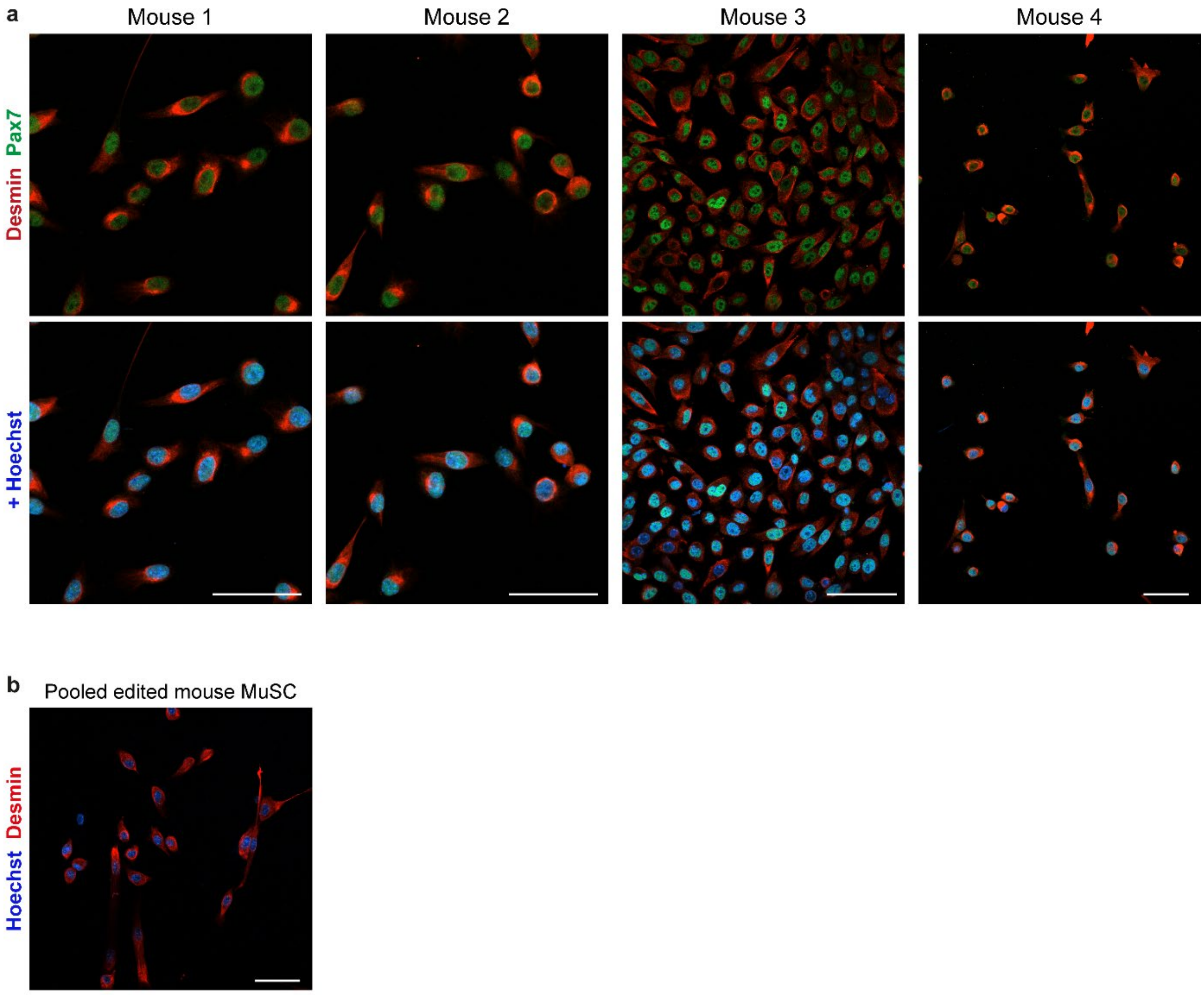


**Supplementary Fig. 11: hEx44mut MuSC remained pure and fit following editing.** **a** Pax7/Desmin immunostaining of MuSC from homozygous hEx44mut mice prior to gene editing. Scale bars: 50 µm. **b** MuSC from homozygous hEx44mut mice remained pure after culture and editing, as shown by the Desmin immunostaining of a subset of the gene edited cell populations that were pooled for transplantation. Scale bar: 50 µm.

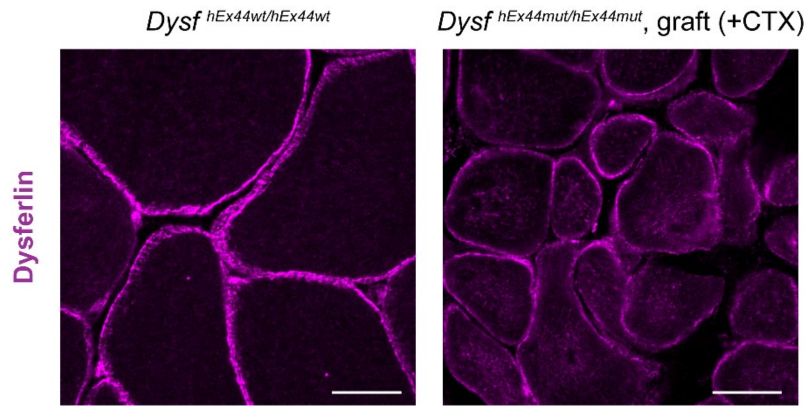


**Supplementary Fig. 12: Re-framed dysferlin shows a similar-to-wild-type localization pattern in donor-derived myofibers.** Dysferlin immunostaining of transversal cryosections from TA muscles of a homozygous hEx44wt mouse (left) and a grafted homozygous hEx44mut mouse (right – the image shows the graft area containing donor-derived, dysferlin-positive fibers). Scale bars: 20 µm.

**Supplementary tables**

**Supplementary Table 1. MuSC populations used in this study**

| **Age at biopsy (y)** | **Gender (m/f)** | **Clinical diagnosis** | **Muscle histology** | **MD mutation** |
|---|---|---|---|---|
| 28 | f | LGMD2B | LGMD2B | *DYSF* exon 44 c.4872-c.4876delinsCCCC |
| 25 | f | LGMD2B | LGMD2B | *DYSF* exon 44 c.4872-c.4876delinsCCCC |
| 24 | f | Suspected myositis | Normal histology | no |
| 23 | f | Suspected myositis | Normal histology | no |
| 21 | m | HyperCKemia of unknown cause | Normal histology | no |

**Supplementary Table 2. gRNA (spacer) sequences**

| **Name** | **Sequence 5' > 3'** (spacer) | **Target** |
|---|---|---|
| DYSFex44mut_gRNA#1 | CTGCACGCTGGACCCCTATT | *DYSF* exon 44 c.4872delinsCCCC |
| DYSFex44mut_gRNA#2 | CAAATAGGGGTCCAGCGTGC | *DYSF* exon 44 c.4872delinsCCCC |
| DYSFex44mut_gRNA#3 | AAATAGGGGTCCAGCGTGCA | *DYSF* exon 44 c.4872delinsCCCC |
| - | ACTTTCCGAATACAGGCTCC | Mouse *Dysf* exon 44 (to generate transgenic mice) |

**Supplementary Table 3. Off-target sites analyzed (OTS)**

| ID | Locus | OTS site (PAM) | Mismatched positions | Chr. | Start | End |
|---|---|---|---|---|---|---|
| **OTS 1** | intergenic:RP11-468 D11.1-RP11-35O7.1 | GAATAGAGGTCCAGCATTCACGG | *.....*........*.*.. | 5 | 2849464 | 2849486 |
| **OTS 2** | exon:SACS | AAAATGGGTTCCAGCCTGCAAGG | ...**...*......*.... | 13 | 23340392 | 23340414 |
| **OTS 3** | intergenic:SPHKAP-AC009410.1 | AACTTGGAGTCCAGCGTTCAAGG | ..*.*..*.........*.. | 2 | 228437695 | 228437717 |
| **OTS 4** | exon:MICALL1 | AAATAGGGTCCCAGGGTCCAGGG | ........**....*..*.. | 22 | 37941104 | 37941126 |
| **OTS 5** | intergenic:RN7SL795P-GTSCR1 | AACTTGGAGTCCAGCGTTCACGG | ..*.*..*.........*.. | 18 | 70549868 | 70549890 |
| **OTS 6** | exon:NLRP1/U7 | AAATTGGGGTTCAGCGTGGGAGG | ....*.....*.......** | 17 | 5514358 | 5514380 |
| **OTS 7** | exon:NUDT18 | GGCCAGGGGTCCAGCGTGCACAG | ****................ | 8 | 22109233 | 22109255 |
| **OTS 8** | intron:HMCN2 | AAGCAGAGGTGCAGCGTGCAGGG | ..**..*...*......... | 9 | 130266781 | 130266803 |
| **OTS 9** | exon:RP11-580I1.2 | CAATAGCAGTCCAGCCTGCATGG | *.....**.......*.... | 15 | 24276480 | 24276502 |
| **OTS 10** | exon:PWRN3 | CAATAGCAGTCCAGTGTGCATGG | *.....**......*..... | 15 | 24442761 | 24442783 |
| **OTS 11** | intron:RP11-305B6.3 | AAATAGGGGGAGAACGTGCATGG | .........***.*...... | 14 | 44149592 | 44149614 |
| **OTS 12** | exon:EDF1 | AACAAGGGGTCCAGCTTGCGGGG | ..**...........*...* | 9 | 136862170 | 136862192 |
| **OTS 13** | intergenic:CNEP1R1-RP11-429P3.3 | AAATGGGACTCCAGCATGCAAGG | ....*..**......*.... | 16 | 50042948 | 50042970 |
| **OTS 14** | intron:NR1H4 | AAATAGGGGAACAGCATCCAAGG | .........**....*.*.. | 12 | 100535509 | 100535531 |
| **OTS 15** | intergenic:RP11-434D2.7-AC008088.4 | TCCTTGGGGTCCAGCGTGCAGGG | ***.*............... | 17 | 20515163 | 20515185 |
| | intergenic:RP11-219A15.4-TNFRSF13B | | ***.*............... | 17 | 16843496 | 16843518 |
| | intergenic:AL353997.3-LGALS9C | | ***.*............... | 17 | 18429224 | 18429246 |

**Supplementary Table 4. List of oligodeoxynucleotides used in the study**

| Oligo name | Sequence 5' > 3' | Purpose |
|---|---|---|
| HE7/DYSF i43 F | CAGGACACAGCCCACATCT | Human *DYSF* exon 44 PCR |
| HE8/DYSF i44 R | CTATGCCCCCATAGACATGC | |
| HE49/Sp_sgRNA_DYSFex44 mut#1_for | CTGCACGCTGGACCCCTATTgtttt | Cloning of spacer sequences into BplI-digested HE_p3.1 (eSpCas91.1) and HE_p4.1 (SpCas9) vector backbones |
| HE50/Sp_sgRNA_DYSFex44 mut#1_rev | AATAGGGGTCCAGCGTGCAGggtgt | |
| HE51/Sp_sgRNA_DYSFex44 mut#2_for | CAAATAGGGGTCCAGCGTGCgtttt | |
| HE52/Sp_sgRNA_DYSFex44 mut#2_rev | GCACGCTGGACCCCTATTTGggtgt | |
| HE53/Sp_sgRNA_DYSFex44 mut#3_for | AAATAGGGGTCCAGCGTGCAgtttt | |
| HE54/Sp_sgRNA_DYSFex44 mut#3_rev | TGCACGCTGGACCCCTATTTggtgt | |
| HE33/humU6prom seq F1 | GGCCTATTTCCCATGATTCC | spacer cloning verification (Sanger) |
| HE85/ssODN-HDR_DYSFex44wt#1 | CCATAGGGAAGAAATCAGTGAGTG ACCAGGATAACTACATCCCCTGCAC GCTGGAGCCCGTATTTGGAAAGTA AATTGGGGCATCTTGGGTCTTGGGG TGGAGGAGCCAGACAGGATAAC | HDR template, human wild-type *DYSF* exon 44 |
| HE307/DYSFex44_OT-ig#1_chr5_Fwd | CTCAGAGCCACAGTGGAAGG | Off-target analysis PCRs (OTS 1-15) |
| HE308/DYSFex44_OT-ig#1_chr5_Rev | TGAAGAGCTGCTGTTGGGAG | |
| HE309/DYSFex44_OT-ig#2_chr16_Fwd | AGTGGATGGATGGGTACAGAC | |
| HE310/DYSFex44_OT-ig#2_chr16_Rev | GCCCACTAAACCACAATTCCT | |
| HE311/DYSFex44_OT-intron:NR1H4_Fwd | ACCTAGTGTTTGCCAGCAGA | |
| HE312/DYSFex44_OT-intron:NR1H4_Rev | AGAAGGGTGTGGTTTGGACA | |
| HE313/DYSFex44_OT-intron:RP11-305B6.3_Fwd | TGCCCAAAACTCCTCATGCT | |
| HE314/DYSFex44_OT-intron:RP11-305B6.3_Rev | CTGGAGAAACAGGCAGGAGC | |
| HE315/DYSFex44_OT-ig#3-5_chr17_Fwd | TGGGAGGTTCCTTGGGTACA | |
| HE316/DYSFex44_OT-ig#3-5_chr17_Rev | CCTGATTCCCATGGCAGGTT | |
| HE317/DYSFex44_OT-ig#6_chr2_Fwd | AGCTGCCAGGGAATATAAAAC | |
| HE318/DYSFex44_OT-ig#6_chr2_Rev | GTTGAAGCCTCCAAGTCTGTG | |
| HE319/DYSFex44_OT-ig#7_chr18_Fwd | TGGGCACCATTTAATCAGTTG | |
| HE320/DYSFex44_OT-ig#7_chr18_Rev | CCTTGAGTAACAACTGCTAGC | |
| HE321/DYSFex44_OT-intron:HMCN2_Fwd | CTTGGAGACCGGAATGGCTC | |

| | | |
|---|---|---|
| HE322/DYSFex44_OT-intron:HMCN2_Rev | GACTCTCTACAGTGGAGCGC | |
| HE323/DYSFex44_OT-exon:NUDT18_Fwd | CGTTGGGTCTTCTGTCCCCA | |
| HE324/DYSFex44_OT-exon:NUDT18_Rev | CAGCCTATCAGCGGCCAGAG | |
| HE325/DYSFex44_OT-exon:PWRN3_Fwd | GGCGTCAGTCTTTGTGCAAT | |
| HE326/DYSFex44_OT-exon:PWRN3_Rev | GGACAGCGATACCTGAGACA | |
| HE327/DYSFex44_OT-exon:NLRP1/U7_Fwd | CTGTTGGCTTGCTCTTGTAGA | |
| HE328/DYSFex44_OT-exon:NLRP1/U7_Rev | GCCTTCCAGCACTAAAGTAATG | |
| HE329/DYSFex44_OT-exon:MICALL1_Fwd | TGCTCCCCTCAGATCAGTCA | |
| HE330/DYSFex44_OT-exon:MICALL1_Rev | CTGGAAGAGCAGAACCCTGG | |
| HE331/DYSFex44_OT-exon:SACS_Fwd | TTCCAGACCAAAGAGCCTGG | |
| HE332/DYSFex44_OT-exon:SACS_Rev | GCTGGTGAACTCTTTGACCCT | |
| HE333/DYSFex44_OT-exon:RP11-580I1.2_Fwd | GGTATGTGTCCACTTGGTTGG | |
| HE334/DYSFex44_OT-exon:RP11-580I1.2_Rev | GACTGGCTGGATCGCATCTA | |
| HE335/DYSFex44_OT-exon:EDF1_Fwd | GAGGAACGCGATGTAGGGAG | |
| HE336/DYSFex44_OT-exon:EDF1_Rev | AGATGTTGAAGAGGGGCAGC | |
| HE14/pMX seq R | CCCAATACGCAAGGAAACAG | Sequencing of mTV_hDYSF_ex44 wt&mut inserts from GeneArt pMX vector |
| HE15/mDysf i43 F2 | GTTGGAAAGGGAGGGAGAAC | |
| HE16/mDysf i44 R2 | TGAAAAGGTCTCTTGGGAAGG | |
| HE17/mDysf i44 F | CCACCAGTTTCTCTTTCTGTCC | |
| HE12/Dysf i43 F | GCTTTCCTTGTGTGCCAGTG | Mouse *Dysf* exon 44 (& hEx44) PCR-RFLP genotyping and genome editing analysis by Sanger |
| HE13/Dysf i44 R | TAAGGGGAAGGTGGGGTGAA | |
| HE223/mDysf e43 F | CAGGAATGCTTGGTCCGTAT | PCR plus Sanger sequencing of the complete hEx44mut and hEx44wt knock-in alleles |
| HE224/mDysf i43 F2 | TTTGGGCTTCTGGAAAAATG | |
| HE225/mDysf e45 F | AGAAGATTGGGGAGACGGTC | |
| HE226/mDysf e45 R | GACCGTCTCCCCAATCTTCT | |
| HE337/mDysf i43 F2 | AATGGGTGAACGGGTGAGAT | Mouse *Dysf* exon 44 (& hEx44) PCR and genome editing analysis by NGS |
| HE338/mDysf i44 R2 | CCCACCCAAGGAATTAGAGACT | |
| SDFp_32 | ATCGTCCGAGCATTTGGCTTA | Mouse *Dysf* exon 43-46, RT-qPCR |
| SDFp_33 | GGTCCAGAGACACAGTAGGTC | |
| oAK26/Gapdh_ex4 F1 | GCATCCTGCACCACCAACTG | Mouse *Gapdh*, RT-qPCR |
| oAK27/Gapdh_ex5 R1 | CATCACGCCACAGCTTTCCA | |

**Supplementary Table 5. Short tandem repeat (STR) analysis of patient hiPSC and blood**

| | Patient 1, blood | Patient 1, hiPSC | Patient 2, blood | Patient 2, hiPSC |
|---|---|---|---|---|
| D10S1248 | 13 | 13 | 13 | 13 |
| Vwa | 14,16 | 14,16 | 14,16 | 14,16 |
| D16S539 | 11,14 | 11,14 | 13,14 | 13,14 |
| D2S1338 | 17,23 | 17,23 | 17,18 | 17,18 |
| D8S1179 | 10,11 | 10,11 | 10,11 | 10,11 |
| D21S11 | 27,30 | 27,30 | 28,31 | 28,31 |
| D19S51 | 13,16 | 13,16 | 12,13 | 12,13 |
| D22S1045 | 15,16 | 15,16 | 15,17 | 15,17 |
| D19S433 | 14 | 14 | 14 | 14 |
| TH01 | 6,7 | 6,7 | 6,7 | 6,7 |
| FGA | 21,23 | 21,23 | 21,23 | 21,23 |
| D2S441 | 14 | 14 | 14 | 14 |
| D3S1358 | 14 | 14 | 17,18 | 17,18 |
| D1S1656 | 12,15* | 12,16* | 12,16 | 12,16 |
| D12S391 | 16,22 | 16,22 | 16,22 | 16,22 |
| SE33 | 17,25.2 | 17,25.2 | 18,25.2 | 18,25.2 |
| Amelogenin | X | X | X | X |

*likely tissue-specific mutation

**Supplementary Table 6. Primary antibodies**

| **Antibody** | **Clone** | **Manufacturer & catalogue number** | **Working dilution** |
|---|---|---|---|
| PAX7 | | Santa Cruz Biotechnology, #sc-81648 | IF - 1:300 |
| PAX7 | P3U1 | Developmental studies hybridoma bank (DSHB). Produced in-house from hybridoma cell line. | IF tissue - undiluted cell culture supernatant |
| Ki-67 | | Thermo Fisher Scientifitc, #RM-9106-S0 | IF - 1:300 |
| MYF5 | C20 | Santa Cruz Biotechnology, #sc-302 | IF - 1:2,000 |
| MYOD | 5.8A | Santa Cruz Biotechnology, #sc-32758 | IF - 1:50 |
| Desmin | | Dako, #M0760 | IF -1:100 |
| Desmin | | Abcam, #ab15200 | IF - 1:2,000 |
| Skeletal Myosin (fast) | MY-32 | Sigma-Aldrich, #M4276 | IF - 1:500 |
| Dysferlin (Hamlet) | | Novocastra, NCL-Hamlet | WB - 1:500 |
| Dysferlin (Romeo) | | Abcam, #ab124684 | IF & WB - 1:150<br>IF tissue - 1:50 |
| Annexin A1 | | Abcam, #ab88865 | IF - 1:80 |
| α-tubulin | | Sigma-Aldrich, #T5168 | WB - 1:2,000 |
| Vinculin | VIN-11-5 | Sigma-Aldrich, #V4505 | WB - 1:200 |
| VCAM1 | | R&D system, #AF643 | FACS - 1:100 |
| PE-anti CD31 | MEC13.3 | DB Pharmingen, #553373 | FACS - 1: 200 |
| PE-anti CD45 | 30F11 | DB Pharmingen, #553081 | FACS - 1: 200 |
| PE-anti Sca1 | E13-161.7 | DB Pharmingen, #553336 | FACS - 1: 200 |
| Laminin | | Sigma-Aldrich, #L9393 | IF tissue - 1:200 |

IF: Immunofluorescence staining

WB: Western blot
